# Improvements to the ARTIC multiplex PCR method for SARS-CoV-2 genome sequencing using nanopore

**DOI:** 10.1101/2020.09.04.283077

**Authors:** John R Tyson, Phillip James, David Stoddart, Natalie Sparks, Arthur Wickenhagen, Grant Hall, Ji Hyun Choi, Hope Lapointe, Kimia Kamelian, Andrew D Smith, Natalie Prystajecky, Ian Goodfellow, Sam J Wilson, Richard Harrigan, Terrance P Snutch, Nicholas J Loman, Joshua Quick

## Abstract

Genome sequencing has been widely deployed to study the evolution of SARS-CoV-2 with more than 90,000 genome sequences uploaded to the GISAID database. We published a method for SARS-CoV-2 genome sequencing (https://www.protocols.io/view/ncov-2019-sequencing-protocol-bbmuik6w) online on January 22, 2020. This approach has rapidly become the most popular method for sequencing SARS-CoV-2 due to its simplicity and cost-effectiveness. Here we present improvements to the original protocol: i) an updated primer scheme with 22 additional primers to improve genome coverage, ii) a streamlined library preparation workflow which improves demultiplexing rate for up to 96 samples and reduces hands-on time by several hours and iii) cost savings which bring the reagent cost down to £10 per sample making it practical for individual labs to sequence thousands of SARS-CoV-2 genomes to support national and international genomic epidemiology efforts.

## Introduction

Real-time, genomic surveillance has become a central component in the management of outbreaks of emerging infectious diseases and has been successfully deployed to understand the transmission patterns of Ebola virus [1,2], Lassa virus [3], Yellow fever virus [4,5] and Influenza [6]. Viral genomes can be used to estimate the rate of viral evolution, monitor circulating lineages as well as identifying any signs of adaptation to hosts, treatments or vaccines. Genome sequencing of SARS-CoV-2 has provided epidemiological insights into the early outbreak [7–9] and a number of studies have adopted our method [8,10–17]. Rapid data sharing allows sequences to be compared across labs or public health authorities and can provide insight into routes of transmission and effectiveness of containment measures[18].

A variety of different approaches have been used to sequence SARS-CoV-2 genomes including RNA metagenomics which was deployed to generate the first SARS-CoV-2 reference genome [7]. Such methods are able to recover the whole genome in an untargeted manner making them ideal for virus discovery but are less suited to genomic surveillance where sequencing large numbers of isolates becomes prohibitively expensive. Hybridisation capture is an approach used to enrich for molecules homologous to the bait sequences although for clinical samples a large number of reads are still required. A third approach is PCR-based target enrichment which produces amplicons that span the viral genome. This method is robust to a large range of input titres [19], is highly specific and scalable to hundreds of genomes[2]. Before SARS-CoV-2, the West African Ebola outbreak of 2013-2016 was the most densely sampled viral outbreak with more genomes generated using amplicon sequencing than all other methods combined [20].

The initial release of the ARTIC SARS-CoV-2 sequencing protocol was released early in the outbreak (Jan 22, 2020) supporting early sequencing efforts in many counties. It consisted of a new primer scheme and laboratory protocol building on a previously published protocol [19] however early adopters experienced some issues. Firstly, amplicon dropouts (regions which are absent regardless of coverage) occurred frequently in specific regions of the genome [21]. Secondly, inefficient barcode ligation reduced the proportion of double-barcoded products resulting in lower sequencing yields. Thirdly, the normalisation procedure to ensure similar numbers of reads from each barcode was laborious and limited the number of samples one person could process.

To address these shortcomings we have modified the primer scheme designed to eliminate amplicon dropouts by adding additional alternate primers (‘alts’) for weak or missing regions. Modifications have also been made to the library preparation protocol to improve demultiplexing and a method to eliminate normalisation has been implemented in a new version of the protocol named ‘GunIt’. Cost and scalability are important factors when aiming to sequence thousands of samples and we have modelled the impact of different strategies to reduce the cost of materials including volume reduction, reagent substitution, flowcell washing, use of 96 native barcodes and nanopore sequencing platform. These findings were used to generate the ‘LoCost’ protocol.

## Results

### Primer schemes

To determine which primer scheme yielded the most complete genome coverage at varying input levels we tested ARTIC V1, V2 and V3 of the primer scheme plus a modification of V1 proposed by Itokawa et al. [21] (see Supplementary Table 4 for definitions). Each scheme was used to amplify a serial dilution of cDNA generated from SARS-CoV-2 cell culture of the strain England/2/2020. The Ct values of the serial dilution ranged from 18 (1e-01) to undetected (1e-07) (Supplementary Table 3). We found that all versions of the primer scheme could generate full coverage at 1e-02 input with 131,072 (2^17^) reads. Drop-outs were apparent where increasing the number of reads analysed gave no further improvements in genome coverage and occurred below 1e-05 for all schemes. V3 was the only scheme to produce maximum coverage between 1e-01 to 1e-04 inputs and produced the highest coverage at 1e-05 while Itokawa produced the highest coverage at 1e-06 (Figure 1). All primer schemes produced partial genome coverage at 1e-06 input (Ct 38) and V1 and V3 generated coverage at the 1e-07 which was negative by qPCR.

Investigating the coefficient of variation (CV) of coverage we found that CV and input exhibit an inverse relationship with higher inputs resulting in lower CV. V3 had the lowest CV between 1e-01 and 1e-05 indicating fewer reads are required to achieve full coverage with that primer scheme (Supplementary Figure 1).

**Figure 1.**
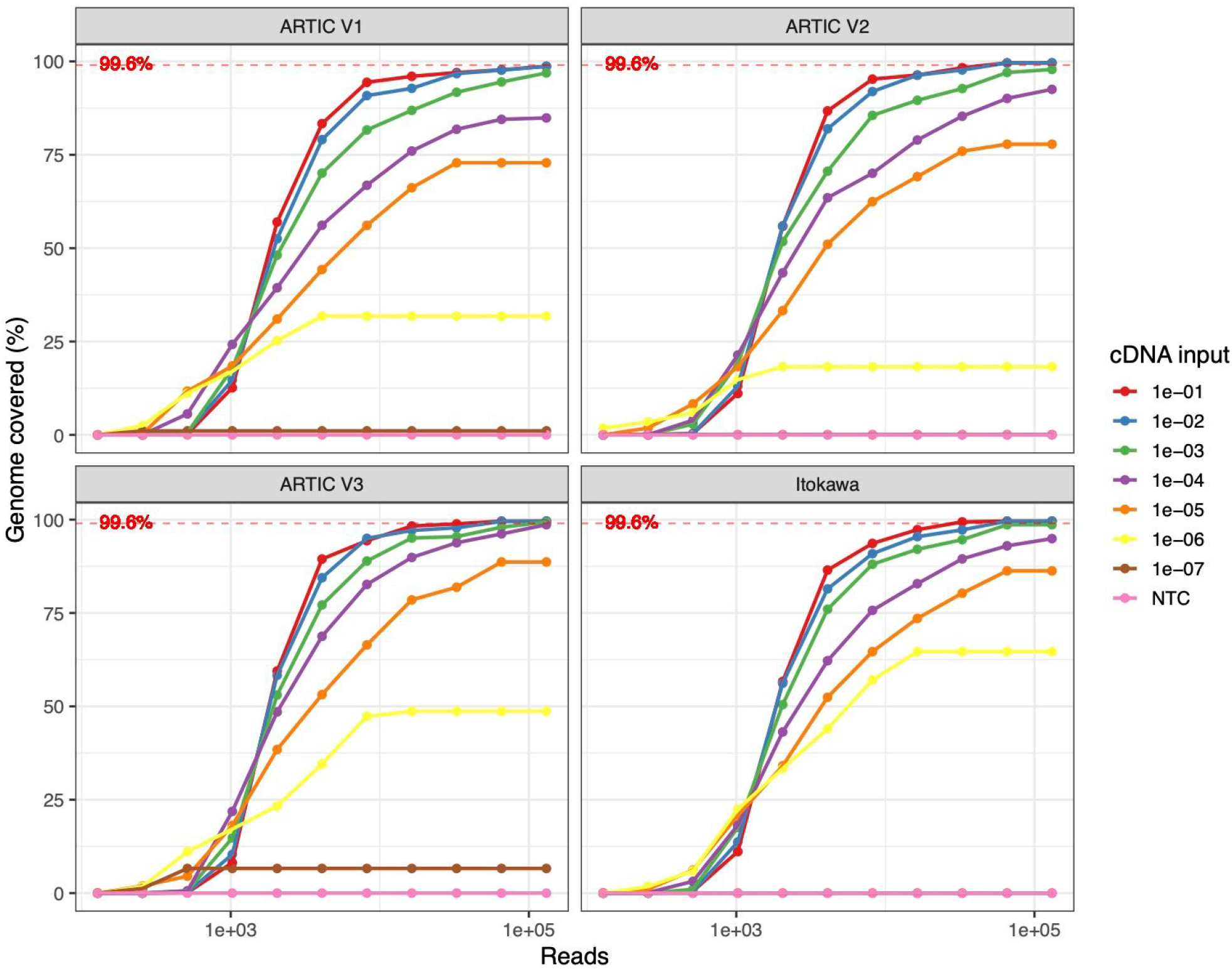
Percentage of genome covered at greater than 20x read depth against read number analysed for different primer schemes and cDNA input. Theoretical maximum genome coverage is indicated by the dotted horizontal line.

### Protocol improvements

The library preparation process converts amplicons generated by the multiplex PCR into barcoded sequencing libraries which can be demultiplexed back into sample bins with high accuracy. This can be achieved by enforcing double-ended barcoding but this typically results in over half the data being discarded [22]. We sought to assess whether protocol changes could increase the amount of usable data generated during a run and to investigate the number of correct and incorrect assignments made.

To do this we designed an experiment in which the first 24 amplicons from the ARTIC V3 primer scheme were barcoded individually so a correctly or incorrectly demultiplexed read could be identified by comparing the mapping position to the barcode assignment. This was repeated for three versions of the sequencing protocol (Baseline, GunIt and LoCost).

All protocols produced a similar number of reads (~5M) in a 12-hour run (Supplementary Table 1). The rate of demultiplexing with double-ended barcoding was 42.15% (Baseline), 67.1% (GunIt) and 74.5% (LoCost) while the rate of incorrectly assigned reads was 0.070% (Baseline), 0.010% (GunIt) and 0.002% (LoCost) using porechop (https://github.com/artic-network/Porechop) with the recommended settings (Table 1). Misassigned reads were reduced 35x using the LoCost protocol with respect to Baseline for the same demultiplexer while increasing the demultiplexing rate by 32.4% (Supplementary Table 2). The lowest rate of misassignment achieved (0.002%) is equivalent to 10 reads per barcode misclassified over a 10M read run.

**Table 1.**
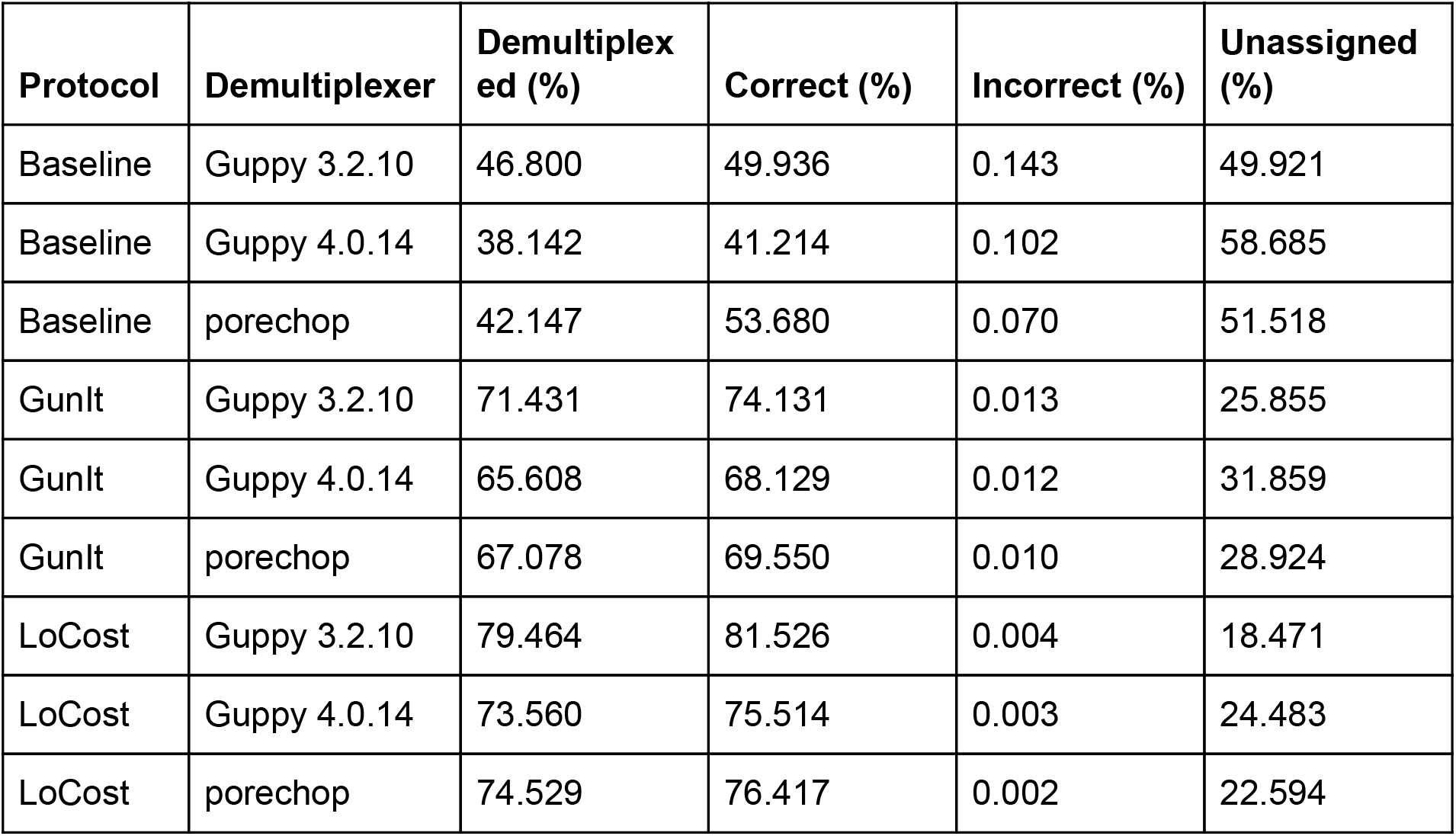
Summary data of demultiplexing outcomes for all protocols for porechop and Guppy using double-ended barcode mode. Full data including absolute read counts shown in Supplementary spreadsheet 1.

| Protocol | Demultiplexer | Demultiplexed (%) | Correct (%) | Incorrect (%) | Unassigned (%) |
| --- | --- | --- | --- | --- | --- |
| Baseline | Guppy 3.2.10 | 46.800 | 49.936 | 0.143 | 49.921 |
| Baseline | Guppy 4.0.14 | 38.142 | 41.214 | 0.102 | 58.685 |
| Baseline | porechop | 42.147 | 53.680 | 0.070 | 51.518 |
| GunIt | Guppy 3.2.10 | 71.431 | 74.131 | 0.013 | 25.855 |
| GunIt | Guppy 4.0.14 | 65.608 | 68.129 | 0.012 | 31.859 |
| GunIt | porechop | 67.078 | 69.550 | 0.010 | 28.924 |
| LoCost | Guppy 3.2.10 | 79.464 | 81.526 | 0.004 | 18.471 |
| LoCost | Guppy 4.0.14 | 73.560 | 75.514 | 0.003 | 24.483 |
| LoCost | porechop | 74.529 | 76.417 | 0.002 | 22.594 |

The results obtained with both porechop and Guppy with the --require_barcodes_both_ends argument were found to be comparable indicating MinKNOW ‘live basecalling’ can be used instead of porechop if this option is enabled. We also investigated single-end barcoding performance and found that although the LoCost protocol reduced misassignments (0.219%) they were still elevated with respect to any condition using double-ended barcoding. Comparing Guppy (3.2.10) which has been in production since the start of the outbreak to the most recent Guppy (4.0.14) we found stringency was increased with respect to the earlier version and produced results equivalent to porechop.

Replacement of the post-PCR clean-up with a 1:10 dilution had no detrimental effect on barcode assignment yet the change removed 1-2 hours of hands-on time from the protocol. The GunIt protocol produced consistently high rates (mean 71.9%) of demultiplexed reads over 8 runs when deployed for clinical sequencing (Supplementary Table 2).

### Run configurations and reagent costs

We sought to investigate the variables that affect material cost including sequencing platform, barcode number, use of flowcell washing, reagents and reaction volumes. We created a model to investigate which changes would have the biggest impact on reducing cost of materials.

There are currently three native barcode expansion packs available providing up to 96 barcodes. While 24 barcodes are sufficient to achieve maximum output on the Flongle, even 96 barcodes are insufficient to achieve maximum output for the MinION/GridION or PromethION flowells without using washing to remove the previous library and reuse the same barcode set (Fig. 2). To assess the feasibility of reusing flowcells with the same barcodes we used the experimental design outlined earlier to determine the carry over from one library to the next when using the flowcell wash kit. By making three libraries each with eight barcodes and loading them sequentially the rate of carry-over after one and two washes could be measured. We determined the carry over to be 0.01% (Supplementary Figure 2) after both the first and second wash.

The modifications included in GunIt (decreasing the volume of barcode ligation reaction and removal of the post-PCR SPRI clean-up) led to a small reduction in cost. Using the model we were able to identify the reagents that contributed significantly to the material cost e.g. NEBNext Ultra II Ligation Module, SuperScript IV Reverse Transcriptase and Native Barcoding Expansion Kit and replace them with alternatives in LoCost. We found the material cost with 24 barcodes on a MinION to be £33.42 using GunIt, £24.91 using LoCost and £16.85 using LoCost with one wash (Figure 3). With 96 barcodes these costs fell dramatically to £18.19 using GunIt, £10.49 using LoCost and £8.47 using LoCost with one wash. A cost of £10.08 can also be achieved on the PromethION with 96 barcodes using the LoCost protocol and two washes.

**Figure 2.**
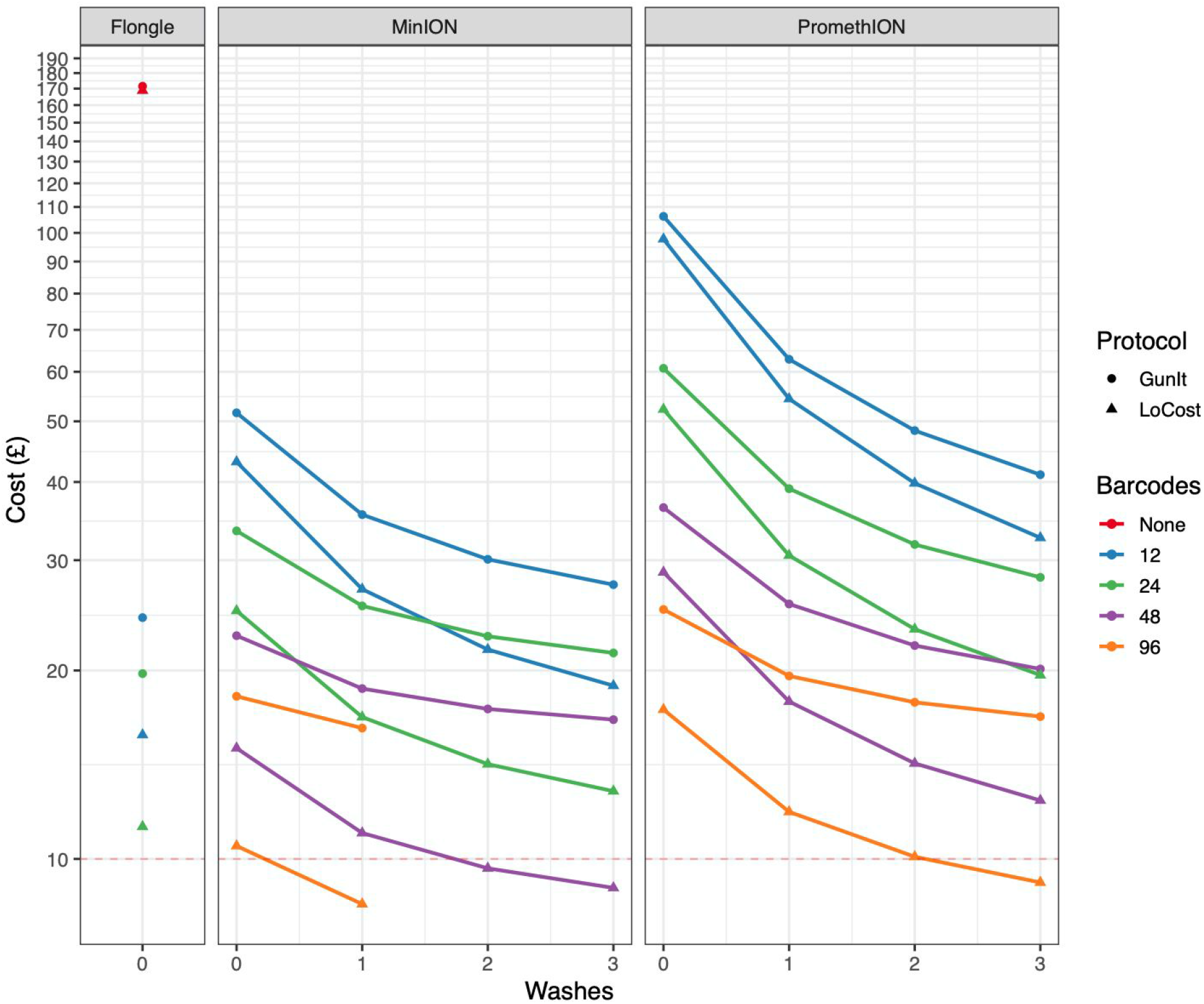
Reagent cost per sample for different barcoding, run configurations and protocol versions. Baseline and GunIt have similar costs so Baseline was removed for clarity. £10 is indicated by the dotted horizontal line.

## Discussion

Here we have addressed several limitations of the ARTIC multiplex PCR method. To eliminate systematic dropouts we introduced the V3 primer scheme which showed the highest genome coverage and lowest CV over a wide range of input cDNA. Modifications to the library preparation protocol have improved the rate of demultiplexing from 42% to 75% while delivering a 35x reduction in the number of incorrectly assigned reads. Further changes to the library preparation protocol reduce the hands-on time by 1-2 hours. Modelling the reagent costs and run configurations allowed us to determine that running 24-48 barcodes on a MinION flowcell is the most cost-effective current run mode with rapid turnaround time. The availability of 96 barcodes allows further cost reduction and makes use of the higher output PromethION instrument cost-effective.

The ARTIC SARS-CoV-2 sequencing protocol has been adopted by many groups around the world due to the low cost, high sensitivity compared to other methods. PCR-based target enrichment produces a high rate of on-target products with an inexpensive and simple laboratory workflow. The main disadvantage is that genome coverage is lost for regions that fail to amplify in the multiplex PCR. Itokawa et al. identified a 10bp overlap between two primers in the initial V1 primer scheme which resulted in the loss of two regions due to heterodimer formation. The scheme was successfully modified but further work developing design tools such as PrimalScheme are needed to prevent this occurring in future.

The V3 primer scheme was found to produce the highest genome coverage with the lowest variance of the primer schemes tested. Reducing the variance reduces the number of reads required, with V3 the requirement is around 100k reads per sample which permits multiplexing up to 96 barcodes on a MinION flowcell. When developing the V3 scheme we spiked 22 alternate primers into the existing pools and found it improved coverage. This led us to hypothesise that the alts provide four possible primer pair combinations per region. This degeneracy could be advantageous when designing future schemes e.g. when spanning regions of variability. Primer mismatches are currently not a major issue affecting coverage due to the relatively low genetic diversity of SARS-CoV-2 genomes circulating globally but new tools such as CoV-GLUE[23] will identify mismatches in diagnostic and sequencing primers/probe sets. Current estimates of SARS-CoV-2 evolutionary rate are 1.04×10^-3^ mutations per site per year which results in 2.6 mutations per month[24]. In future the scheme design is likely to need updating to prevent mismatches in primer sequences.

Itokawa et al. have reported improved overall coverage with their scheme than V3[21]. We believe these differences are explained by slight variation in thermocycler calibration across different sites. Thermocyclers with a lower calibration encourage interactions between primer pairs while those that are high are likely to observe primers with lower Tm dropping out. We have determined that optimal results are achieved when annealing/extension temperature is fine-tuned on any given thermocycler. This remains a minor weakness of the method and further work is needed to reduce this temperature dependence through improved primer design.

Other groups have proposed alternate primer schemes using longer amplicons to improve genome coverage [25,26]. Our primer scheme uses 400 bp amplicons which are compatible with both long and short-read sequencing platforms and offer good performance with degraded or high Ct samples. RNA degradation occurs for a number of reasons including improper storage and issues have been exacerbated during this outbreak due to shortages of lysis buffer and insufficient ultra-low freezer capacity[27]. A scheme length of 400 bp enables efficient amplification when input material is 400 bp or longer increasing the proportion of positive samples that can be amplified compared to longer amplicon schemes. Although not the focus of this study we found the sensitivity of the multiplex PCR with 96 amplicons can equal or even surpass that of qPCR detection which could allow diagnostics by sequencing[28]. In low viral titre samples the absence of one or more diagnostics targets would be called undetermined or negative whereas the complete failure of all 96 amplicon regions across the whole of the SARS-CoV-2 genome is unlikely. The drawback of this approach is that it is more costly and harder to control contamination as unlike RT-qPCR the PCR plates must be unsealed to prepare sequencing libraries. Incorporating dUTP into cDNA/PCR products and subsequent UDG digestion is one way to mitigate these concerns.

Modifications to the library preparation protocol were successful in increasing demultiplexing rate. Although we only have a single dataset for each protocol the rate of demultiplexing was consistently high over 8 runs of clinical samples using GunIt. We found the read misassignment rate reduced using LoCost which reduces the likelihood of incorrect genome sequence resulting from demultiplexing errors. We hypothesise that this improvement was a result of reducing free-barcode in the AMII ligation which improves read-detection for short amplicon sequencing. The protocol changes introduced here eliminate this problem for up to 96 barcodes and can be applied to other applications that use native barcoded sequencing libraries. Removal of the normalisation step with a simple dilution produced no negative effects and removed significant hands-on time from the protocol. It was found that the PCR reaction itself acts as a normalisation step due to the concentration plateauing at ~100 ng/ul by cycle 35 allowing a nominal input to be achieved using a fixed volume of PCR reaction mixture as input to the end-preparation reaction.

Although SARS-CoV-2 genome sequencing using amplicon sequencing is cost-effective compared to other methods our LoCost protocol further reduced material cost to at or below a £10 per sample. The most cost-effective run configurations are 12 barcodes on a Flongle or 96 barcodes on a MinION flowell although cost is only one consideration as it would take significantly longer to generate the same coverage using a Flongle than a MinION flowcell. The MinION can generate at least 20M 400 bp reads on a native barcoded library and therefore requires 96-barcodes or flowcell washing to utilise the yield. The recently released Native Barcode Expansion 96 (EXP-NBD196) kit dramatically reduces the per sample material cost from £24.91 to £10.49.

Carry-over from a previous library after washing was found to be 0.01%, sufficiently low enough that if used judiciously washing can be used to further reduce costs by loading the same barcode set again. We expect this to be less relevant with the 96 barcode expansion kit as it will allow the freedom to run up to 96 barcodes simultaneously or 24-48 barcodes loaded sequentially with washing without reusing the same barcode sequences. This would be the preferred configuration if time to answer, or only a smaller number of daily samples sets are available.

The single largest cost reductions are achieved with a single wash with further washes resulting in incrementally smaller gains. This is because the flowcell becomes an increasingly smaller contributor to the overall cost the more washes are performed. One limitation of this model is that it does not take into account any loss of active pores associated with washing which is why we limited the maximum number of washes to three per flowcell. Cost reductions in LoCost were achieved by identifying alternatives to the most expensive components in use and replacing them with cheaper alternatives or reducing the volume used.

In summary, we have further developed our popular ARTIC SARS-CoV-2 sequencing method improving genome quality, workflow and cost. The V3 primer scheme gives the highest genome coverage and lowest CV of the schemes tested while improvements to the library preparation protocol have improved the rate and accuracy of demultiplexing while also reducing hands-on time. Cost modelling shows that the MinION is the most cost-effective solution to sequence 24 samples rapidly while the 96 native barcode kit provides the highest throughput solution for the MinION and PromethION. The LoCost library preparation protocol reduces the reagent cost to ~£10 per sample which will support the ongoing population scale sequencing efforts underway to monitor the spread of SARS-CoV-2 in the United Kingdom, Canada and elsewhere.

## Methods

### Primer scheme testing study

#### Viral cell culture

Vero E6 cells were maintained in Dulbecco’s modified Eagle’s medium (DMEM) supplemented with 10% fetal calf serum (FCS) and 10 μg / ml gentamicin. Infections were carried out in DMEM containing 2% FCS and gentamicin. All work with infectious virus was carried out in a microbiological safety cabinet within a containment level 3 (CL3) suite observing local biosafety regulations.

The clinical isolate SARS-CoV2 England/2/2020 (received from PHE) was propagated on Vero E6 cells. Initial virus stock from PHE was used to inoculate T25 flasks with 750,000 Vero E6 cells at an estimated MOI of 0.01. After 96-hours clear CPE was observed, and the virus stock was harvested. The supernatant was centrifuge clarified and aliquoted as passage 2 (P2) master stock for further propagations. Subsequent P2 virus stock was used to generate a passage 3 (P3) virus stock by inoculating Vero E6 cells. After 72-hours the virus was harvested by centrifuge clarification of the supernatant and stored at −80C. Virus titre was determined by standard plaque assay. SARS-CoV2 P3 with 1.75×10^5 PFU / mL was subsequently used for experiments.

To obtain an RNA sample of SARS-CoV2, a 6-well plate with 4×10^5 Vero E6 cells / well was infected with 500 PFU / well of SARS-CoV2 P3 in a 1 ml volume. After 1 hour virus absorption the inoculum was replaced with 3 ml fresh DMEM + 2% FCS and gentamicin. 3-days post-infection clear signs of CPE were visible, cells were washed once with PBS and 1 mL Trizol was added to each well. After 5 minutes incubation at room-temperature samples were transferred to screw-cap tubes and heat-inactivated at 80°C for 10 minutes before export from the containment facilities.

#### RNA extraction

RNA was extracted using a hybrid Trizol-RNeasy protocol. In short, 200 uL chloroform was added to each sample of 1 ml Trizol. Tubes were mixed vigorously by hand and centrifuged at 12,000 xG for 15 minutes at 4°C. The aqueous phase was removed and mixed with ½ volume of 100% ethanol before adding to a column of the RNeasy kit (Qiagen). After addition to the column, the standard protocol of the RNeasy kit was followed. Samples were eluted twice in 50 ul of dH_2_O and concentrations and purity were checked on a Nanodrop.

#### cDNA generation

RNA samples were directly used for first-strand synthesis using the SuperScript III kit (ThermoFisher) and random hexamers. Each RNA sample was used for multiple cDNA reactions. In brief, 11 μL RNA were mixed with 1 μL random hexamers (50 μM, Invitrogen) and 1 μL dNTPs. The mixture was incubated for 10 minutes at 65°C and placed directly on ice. After 2-3 minutes, 8 μL enzyme mix containing 4 μL 5x First Strand Buffer, 1 μL 0.1M DTT, 1 μL RNase OUT (Invitrogen) and 2 μL SuperScript III was added to the samples. The reactions were placed in a thermocycler and incubated 5 minutes at 25°C, followed by 1 hour at 50°C and 5 minutes at 90°C before cooling to 4°C. cDNA reactions from the same RNA sample were pooled together and stored at −20°C.

#### Generation of dilution series

A serial dilution of the cDNA was performed from 1e-01 to 1e-07 by taking 5uL of cDNA diluting it in 45uL nuclease-free water before mixing by pipetting and repeating the process 7 times.

#### qPCR of control material

qPCR was performed using the 2019-nCoV CDC qPCR Probe Assay (https://www.cdc.gov/coronavirus/2019-ncov/lab/rt-pcr-panel-primer-probes.html). 25 μL reactions for each of the targets were set up as follows; 1x GoTaq Probe Mastermix (Promega), 500nM each forward and reverse primer, 125nM probe (IDT), nuclease-free water and 2.5 μL cDNA. This was run on a LightCycler 96 (Roche) using the following program; 95°C for 120 secs followed by 45 cycles of 95°C for 10 secs and 55°C for 30 secs and configured for detection of FAM fluorescence. Data was analysed using the relative quantification template in the instrument software.

#### SARS-CoV-2 scheme design

A multiplex PCR primer scheme was designed using primalscheme commit https://github.com/aresti/primalscheme/commit/4fe78b9a1515949aa2b98ff7e69ec23f0aedc3 8a using the SARS-CoV-2 reference genome MN908947.3 specifying an amplicon length of 400 bp and a minimum overlap of 0 bp.

#### Library preparation and sequencing

Primer pools were prepared for V1, V2, V3 and Itokawa (all 15nM each primer pre-pooled to 10 μM). The rest of the library preparation followed the GunIt protocol. A 12 hour run was initiated using MinKNOW (19.12.6) with high-accuracy basecalling using Guppy (3.2.10). The MinKNOW config was modified using the following command;

`/opt/ont/minknow/bin/config editor --conf application --filename`

`/opt/ont/minknow/conf/app_conf --set`

`guppy.extra_arguments=“--require_barcodes_both_ends”`

#### Analysis

Reads were collected and filtered using the field bioinformatics pipeline (https://github.com/artic-network/fieldbioinformatics) `guppyplex` command with the parameters `--skip-quality-check --min-length 400 --max-length 700`. Reads were subsampled using `seqtk` (https://github.com/lh3/seqtk) in batches doubling in number from 128 to 131072. Reads were processed using the minion command using the parameters `--normalise 0`. Coverage was calculated from the primer trimmed BAM files using `samtools depth` command (https://github.com/samtools/). CV was calculated using scipy.stats.variation.

### Barcode assignment study

#### Amplicon generation

Primers pairs from the ARTIC V3 primer scheme were used to amplify amplicons 1-24 individually from SARS-CoV-2 cDNA. Singleplex PCR conditions were as follows; 1x Q5 Reaction Buffer, 200 μM dNTPs, 0.2 μM forward and reverse primer, 0.02 U/μl Q5 High-Fidelity DNA Polymerase, 2.5 uL SARS-CoV-2 cDNA (see above). PCR cycling conditions were; 98°C for 30 secs followed by 35 cycles of 98°C for 15 secs, 65°C for 5 mins.

#### Library preparation and sequencing

Sequencing libraries were prepared using the Baseline (https://www.protocols.io/view/ncov-2019-sequencing-protocol-bbmuik6w), GunIT (https://www.protocols.io/view/ncov-2019-sequencing-protocol-v2-bdp7i5rn) or LoCost (https://www.protocols.io/view/ncov-2019-sequencing-protocol-v3-locost-bh42j8ye). Up to 15 ng library was loaded onto the flowcell and a 12-hour run performed with high-accuracy basecalling (Guppy 3.2.10) but demultiplexing turned off. Changes included in these updated protocols were;

GunIt

- Post-PCR SPRI clean-up replaced with a 1:10 dilution in nuclease-free water.
- Native barcode ligation reaction volume reduced to 20 uL.
- Post-ligation SPRI ratio reduced from 1x to 0.4x and short fragment buffer (SFB) washes added.

LoCost

- Substitution of SuperScript IV for LunaScript RT SuperMix and reaction volume reduced to 10 uL.
- Substitution of Ultra II Ligation Module for Blunt/TA Ligase Master Mix and reaction volume reduced to 10 μL.
- Native barcode ligation reaction volume reduced to 10 uL.
- SFB wash volume reduced.

#### Demultiplexing

Demultiplexing was performed using porechop (https://github.com/artic-network/Porechop), Guppy 3.2.10 as released with MinKNOW (19.12.6) or Guppy 4.0.14 (https://mirror.oxfordnanoportal.com/software/analysis/ont-guppy_4.0.14_linux64.tar.gz) with the following commands;

`guppy barcoder [--require_barcodes_both_ends] -i fastq_pass -s output_dir --arrangements_files “barcode_arrs_nb12.cfg barcode_arrs_nb24.cfg”`

porechop --verbosity 2 --untrimmed -i fastq_pass -b output_dir --native_barcodes --discard_middle [--require_two_barcodes] --barcode_threshold 80 --check_reads 10000 --barcode_diff 5 > demultiplexreport.txt

#### Analysis

Reads for each protocol were aligned to the MN908947.3 reference genome using minimap2 [29]. The BAM file was filtered for primary alignments using the bitwise flag 0x904. The ground truth was assigned for each readset using `bedtools intersect` [30] with the options `-F 1.0 -f 0.8` to remove low confidence assignments. dplyr (RStudio) was used to join the `barcoding_summary.txt` with the BED file produced by bedtools using the read ID. Reads where the called barcode matched the ground truth was designated correct, where it matched a different barcode it was designated incorrect and where a barcode was not assigned it was designated unassigned.

### Library carry-over study

#### Amplicon generation

Primers in Supplementary Table 1 were used to amplify 21 discrete amplicons from *Escherichia coli* DH5α genomic DNA. Reactions were carried out in uniplex 50 μL reactions using 0.5 μM of each primer pair, 30ng of *E. coli* DH5α genomic DNA, 200μM dNTPs, 1X Q5 Reaction Buffer (NEB) and 0.02 U/μL Q5 Hot Start High-Fidelity DNA Polymerase. PCR cycling conditions were; 98°C for 30 seconds followed by 30 cycles of 98°C for 15 seconds, 61°C for 10 seconds and 72°C for 30 seconds. The reaction concluded with a final extension of 72°C for 5 minutes. The amplicons underwent 0.7X SPRI (Aline Biosciences) purification with 2 × 80% ethanol washes (200 μL) and eluted in 30 μL of NFW.

#### Library preparation and sequencing

Amplicons were quantified using the Qubit dsDNA HS kit (Thermo) and 50 ng (~0.08pmol) was taken forward to an end-prep reaction containing 0.75 μL of Ultra II End-prep enzyme mix and 1.75 μL 1X Ultra II End-prep buffer in a total volume of 15 μL. The reaction was carried out at 25°C for 15 minutes followed by 75°C for 10 minutes in individual reactions. The end-prepped amplicons underwent 0.7X SPRI (Aline Biosciences) purification with 2 × 80% ethanol washes (200μL) and eluted in 30 μL of NFW. 10 ng (~0.016 pmol) of end-prepped amplicons were taken forward to a native barcode ligation step containing 2.5 μL of one of 24 native barcodes (Oxford Nanopore Technologies), 0.5 μL of Ultra II Ligation Module Enhancer and 10 μL of Ultra II Ligation Module master mix (NEB) made up of a total of 20 μL with NFW. Ligation was carried out at room temperature (25°C) for 20 minutes prior to heat inactivation at 75°C for 10 minutes. Barcoded amplicons were pooled into three pools of 8 and a 0.4X SPRI purification was carried out with 2 × 250 μL of SFB (ONT) washes and a final 200 μL 80% ethanol wash. Amplicons were stored at −20°C until required.

For each library 30 μL of purified pooled barcoded amplicons (~35 ng) was taken forward to a 50 μL adapter ligation step where 5 μL of AMII (ONT), 5 μL of Quick T4 Ligase (NEB) in 1X Quick Ligation buffer (NEB). Ligation was carried out for 20 min at room temp (25°C) and then underwent a 1X SPRI (Aline Biosciences) purification with 2 × 250 μL SFB washes and eluted in 15 μL EB (ONT). Each library was then quantified and approximately 23.8 ng (~35 fmol) was loaded onto the library. The first library was loaded onto a FLO-MIN106 9.4.1 (ONT) and sequenced until all amplicons had 100,000 reads. The experiment was ended and the flowcell washed with the Flowcell Wash Kit EXP-WSH003 (ONT) as per manufacturer instructions and underwent a 30 minute incubation before being primed for the next library. This was repeated for the second and third library. Runs were refuelled with FB if the translocation speed dropped outside of the recommended range.

#### Analysis

Read demultiplexing was enabled in MinKNOW with --require_barcodes_both_ends configured. The kit SQK-LSK109 and barcoding expansion packs EXP-NBD104 and EXP-NBD114 were specified for all three libraries. Pass reads were counted from each the FASTQ files and then analyzed in Prism 8.0. Percentages were calculated as the sum of specified reads in each library. Other run stats were gathered from the experiment summary file. Reads were counted using: awk `‘{s++}END{print s/4}’`

`fastq pass/barcode{}/*.fastq >> reads.csv`

### Cost model

Reagent prices were taken from Science Warehouse available to the University of Birmingham on 1st April 2020. Flowcell costs were based on individual pricing for Flongle and for flowcells packs of 48 (MinION/GridION) and 12 (PromethION). The price for each series was calculated using an Excel spreadsheet until the estimated maximum flowcell output of 2.4M (Flongle), 19.2M (MinION/GridION) or 96M (PromethION) was reached or the number of washes became impractical (>3). ‘Baseline’ indicates the initial protocol release (dx.doi.org/10.17504/protocols.io.bbmuik6w), ‘GunIt’ refers to the release (dx.doi.org/10.17504/protocols.io.bdp7i5rn) and ‘LoCost’ includes LunaScript RT SuperMix (NEB) instead of SuperScript IV (Thermo) and half volume native barcoding reactions using Blunt/TA Ligase Master Mix (NEB) instead of NEBNext Ultra II Ligation Module (NEB).

## Author contributions (CRediT taxonomy)

Conceptualization, JQ and NJL; Methodology, JRT, PJ, DS and JQ; Software, JQ, AS and NJL;

Formal Analysis, JQ, JRT, PJ, DS and GH;

Investigation, JQ, JRT, NS, JC, HL, KK, AW and GH; Resources, NJL, NP, RH, SW, TPS;

Writing – Original Draft, JQ;

Writing – Review & Editing, JQ, JRT and NJL;

Visualization - JQ, PJ and GH;

Supervision, JQ, NJL, TPS, RH, NP, SJW and IG; Funding Acquisition, JQ, NJL, NP, RH and TPS.

## Acknowledgements

We would like to acknowledge the groups internationally that used the protocol on release and provided useful feedback and datasets. We thank PHE for allowing access to the England/2/202 strain for development work.

This work was supported by the ARTIC project Wellcome Trust Collaborative Award. The MRC-University of Glasgow CVR is supported by MRC grant MC PC 19026. JQ is funded by a UKRI Future Leaders Fellowship. JRT and TPS were supported by the Canadian Institutes of Health Research (#10677), the Canadian Epigenetics, Environment and Health Research Consortium Network, and Genome Canada through the Canadian COVID-19 Genome Network.

## Data availability

Raw read data are available for download, barcode-sample mapping in Supplementary Table 5.

### Primer scheme testing study

https://artic.s3.climb.ac.uk/GunIt_nb_serial_V2V1.tgz

https://artic.s3.climb.ac.uk/GunIt_nb_serial_V3It.tgz

### Barcode assignment study

https://artic.s3.climb.ac.uk/Baseline_NB_Single.tgz

https://artic.s3.climb.ac.uk/GunIt_NB_Single_2.tgz

https://artic.s3.climb.ac.uk/LoCost_NB_Single.tgz

