## Supplementary information for "Improvements to the ARTIC multiplex PCR method for SARS-CoV-2 genome sequencing using nanopore"

| **Start position** | **End position** | **Forward/reverse primer** | **Reverse primer** |
| --- | --- | --- | --- |
| 2279 | 3224 | CGAAGGCATGAGTTTCTCCG | TGTCGTAATGAATGCTGCCG |
| 281973 | 282896 | GCGACTCCATAAACGGGTTC | CGAGCAGCAGGGTATTGAAC |
| 460512 | 461372 | TGGTCTGGAGCGTGAAATCT | CATCCCCTTACAACACCCCT |
| 518953 | 519838 | CGATCAACACGCCGGTAAAT | CACGTTTGACAACCAGCTCA |
| 742504 | 743437 | GCAGCGCAACGACATAGTAA | CGGGCTGAATGAATGGTCTG |
| 876707 | 877643 | AATTAAACCCGGCTACCCCA | ACAAGGTGTGAAACTGACGC |
| 1028462 | 1029325 | TGGATGGCTGTTCAGGATGT | CCGCCGCTACGTTGAAAATA |
| 1055605 | 1056589 | CCGATCGAGTGCACTTTCAG | CTACCGAGCAAAATGGCCTC |
| 1390082 | 1391077 | GCCTAATTCGCTGTGTCTGG | TGACGACCTGTACGATCGAG |
| 1780903 | 1781783 | TTATGTTGCTCCGCGTCATG | CGAACGCCTGGGATAAACTG |
| 1968363 | 1969238 | CTTAATCAAGTCACGCGCGA | GCTGACCATAAACGCGACAT |
| 2126280 | 2127172 | CCGCTTCCAGTGAGATTTCG | TTTATTGCTGGTCATCGCGG |
| 2195755 | 2196680 | GCGTGCTGATGACTTACCTG | CTTCGCTTGCTCCACATACC |
| 2269263 | 2270216 | CACGAAAGATGCCTCACGAG | CGCCTTTCCAGTCAGCATAC |
| 2593940 | 2594791 | CGGACCAGTAACATCTTGCG | ACGTTATGGCAAAGCGATCC |
| 2740934 | 2741800 | TTCCAGAGCAGTTTCCGGAT | ATGCGACCGTCATCAATGTG |
| 3239321 | 3240273 | TGCGTGCGATGTATTTCCTG | CAGGGTAGAAGGGAAGGCAA |
| 3537686 | 3538630 | AGACCTCTACCTGCCCTACA | TGACGATTACTTTCTGCGCG |
| 3920030 | 3920979 | TTTTCGCGAGCTGTTGTTCA | GGTCTGTACGTGATGTTCGC |
| 4016553 | 4017460 | GTTAAGCTGGCATCTCACCG | GCGACCATCTCTTCACGTTC |
| 4293990 | 4294882 | TGGCTTCCGATGCGTTTATG | GCTTTGTACCGCTTCCGTAG |
| 4226892 | 4227804 | GATCAACGTCAGCCACCATC | ACGCCGGTTCGAACAATATG |
| 4546199 | 4547124 | CAACGTGAAGGGCATACTCG | AACCACCGCGATAATTCAGC |
| 4594375 | 4595263 | AAGAACATCCCCAGTCCCAG | CTACTGCGTGACAACATCGG |

Supplementary Table 1. Coordinates against NC_000913.3 and primer sequences for E. coli amplicons used in the wash experiment.

| **Run date** | **Barcodes Samples** | **Total pass reads (16hr run time)** | **% double barcoded pass reads (>=q7)** |
| --- | --- | --- | --- |
| 22nd March 2020 | 24 | 10.4M | 76.2 |
| 24th March 2020 | 24 | 8.2M | 70.8 |
| 24th March 2020 | 21 | 10.8M | 60.4 |
| 1st April 2020 | 24 | 12.0M | 70.7 |
| 1st April 2020 | 24 | 10.5M | 71.0 |
| 3rd April 2020 | 24 | 11.2M | 76.2 |
| 3rd April 2020 | 24 | 10.2M | 72.5 |
| 3rd April 2020 | 24 | 10.3M | 77.7 |
| Average (mean) | | 10.5M | 71.9 |

Supplementary Table 2. BCCDC barcoding data for early GunIt runs. (Double end barcoded pass (>=q7) reads expressed as a percentage of total pass reads (barcode assigned + unclassified).

| **Dilution** | **N1 (Ct)** | **N2 (Ct)** |
| --- | --- | --- |
| 1e-01 | 17.18 | 18.03 |
| 1e-02 | 20.46 | 21.46 |
| 1e-03 | 24.73 | 25.76 |
| 1e-04 | 29.84 | 30.74 |
| 1e-05 | 33.3 | 34.33 |
| 1e-06 | 38.26 | 37.36 |
| 1e-07 | - | - |
| NTC | - | - |

Supplementary Table 3. qPCR cycle threshold (Ct) values for dilution series used in primer scheme comparison study.

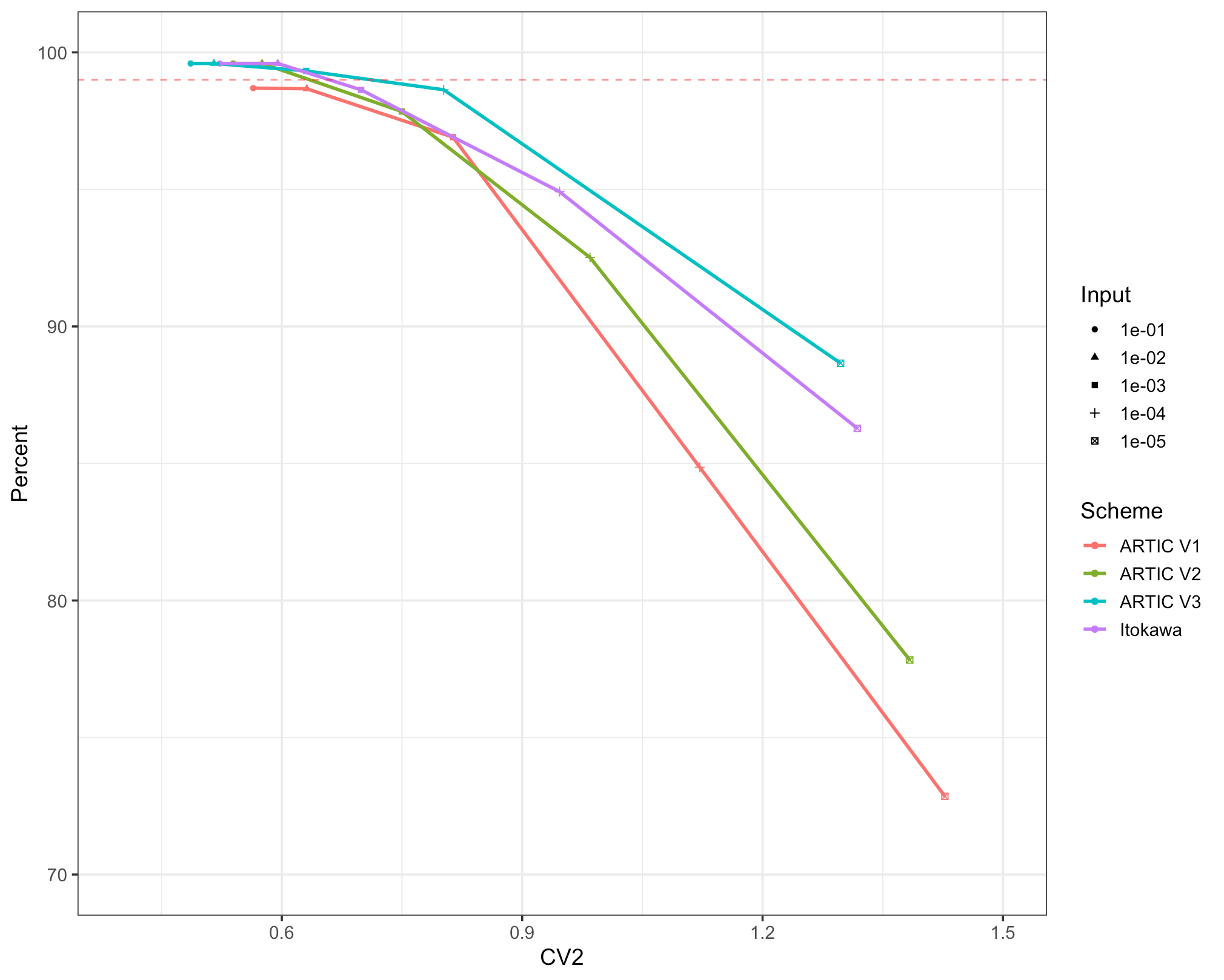

Supplementary Figure 1. Genome coverage vs coefficient of variation (CV) of coverage for different primer schemes and inputs. Theoretical maximum genome coverage is indicated by the dotted horizontal line.

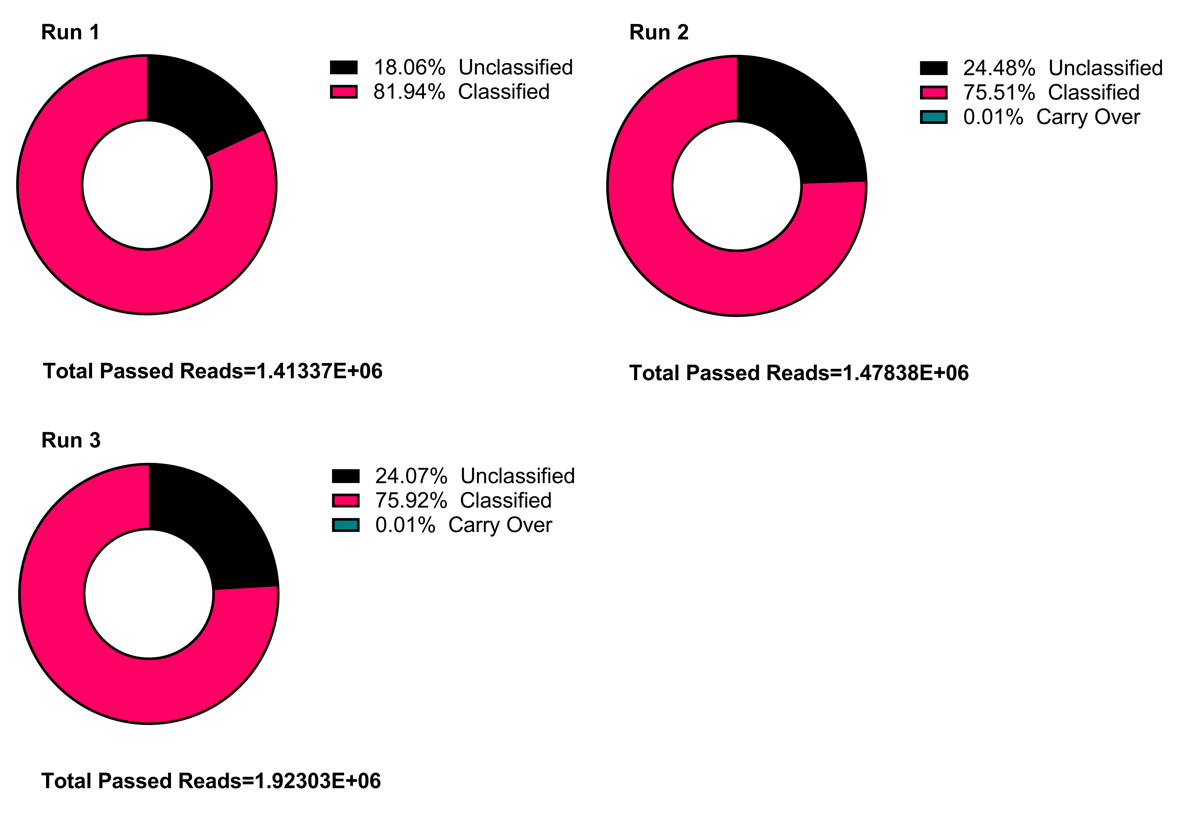

Supplementary Figure 2. Total reads and percentage of unassigned, demultiplexed, and carry over reads for three sequential runs with washes between them.

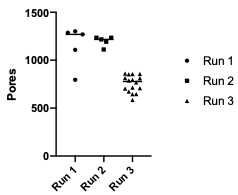

Supplementary Figure 3. Available pore number from mux scans by run.

| **Name** | **Definition** |
| --- | --- |
| ARTIC V1, V2, V3, Itokawa et al. | SARS-CoV-2 primer schemes |
| Baseline/GunIt/LoCost | SARS-CoV-2 sequencing protocol versions on protocols.io |
| PrimalScheme | The generic name for a primer scheme |
| PrimalSeq | The generic name for the sequencing method using a PrimalScheme |
| primalscheme | Software (CLI) |

Supplementary Table 4. Definitions for terms used in this manuscript

| **Run name** | **Protocol** | **Barcode-sample mapping** |
| --- | --- | --- |
| GunIt_nb_serial_V2V1 | GunIt | NB01=10-1, V2 scheme  NB02=10-2, V2 scheme  NB03=10-3, V2 scheme  NB04=10-4, V2 scheme  NB05=10-5, V2 scheme  NB06=10-6, V2 scheme  NB07=10-7, V2 scheme  NB08=NTC, V2 scheme  NB09=10-1, V1 scheme  NB10=10-2, V1 scheme  NB11=10-3, V1 scheme  NB12=10-4, V1 scheme  NB13=10-5, V1 scheme  NB14=10-6, V1 scheme  NB15=10-7, V1 scheme  NB16=NTC, V1 scheme |
| GunIt_nb_serial_V3It | GunIt | NB01=10-1, V3 scheme  NB02=10-2, V3 scheme  NB03=10-3, V3 scheme  NB04=10-4, V3 scheme  NB05=10-5, V3 scheme  NB06=10-6, V3 scheme  NB07=10-7, V3 scheme  NB08=NTC, V3 scheme  NB09=10-1, Itokawa scheme  NB10=10-2, Itokawa scheme  NB11=10-3, Itokawa scheme  NB12=10-4, Itokawa scheme  NB13=10-5, Itokawa scheme  NB14=10-6, Itokawa scheme  NB15=10-7, Itokawa scheme  NB16=NTC, Itokawa scheme |
| Baseline_NB_Single | Baseline | NB01=Amplicon 1  NB02=Amplicon 2  NB03=Amplicon 3  NB04=Amplicon 4  NB05=Amplicon 5  NB06=Amplicon 6  NB07=Amplicon 7  NB08=Amplicon 8  NB09=Amplicon 9  NB10=Amplicon 10  NB11=Amplicon 11  NB12=Amplicon 12  NB13=Amplicon 13  NB14=Amplicon 14  NB15=Amplicon 15  NB16=Amplicon 16  NB17=Amplicon 17  NB18=Amplicon 18  NB19=Amplicon 19  NB20=Amplicon 20  NB21=Amplicon 21  NB22=Amplicon 22  NB23=Amplicon 23  NB24=Amplicon 24 |
| GunIt_NB_Single_2 | GunIt | As above |
| LoCost_NB_Single | LoCost | As above |

Supplementary Table 5. Barcode-sample mapping for sequencing runs.
